# The first chromosome-scale genome assembly of *Blumeria graminis* f. sp. *avenae* provides insights into genome evolution and host specialization

**DOI:** 10.64898/2026.08.28.747853

**Authors:** Yi Ding, Peng Zhang, Tomasz Ociepa, Aleksandra Nucia, Haixia Guan, Krzysztof Kowalczyk, Robert F. Park, Sylwia Okoń

## Abstract

*Blumeria graminis* f. sp. *avenae* (Bga), the causal agent of oat powdery mildew, is one of the most host-specialized members of the *B. graminis* species complex. Despite its agricultural importance, the lack of a high-quality reference genome has limited studies of host specialization, virulence evolution and comparative genomics in this pathogen. Here, we generated the first chromosome-scale genome assembly of Bga using an integrative approach combining long-and short-read sequencing, Hi-C scaffolding and transcriptome data. The Bga genome exhibits hallmark features of powdery mildew fungi, including extensive repeat content and low gene density. Comparative analyses revealed that genome expansion is primarily associated with historical transposable element proliferation rather than recent transpositional activity. Genome organization is consistent with a functionally stratified “one-speed” model, in which genes associated with pathogenicity, including predicted effectors and infection-responsive genes, are preferentially located in transposable element-rich regions characterized by reduced synteny conservation and extended intergenic spaces. In contrast, conserved genes are concentrated in compact genomic regions and maintain strong syntenic conservation across cereal-infecting formae speciales. Hi-C analyses demonstrated a highly structured chromatin architecture and revealed genome organization patterns associated with infection-related gene expression. Comparative genomic analyses indicated that host specialization in Bga is driven by localized diversification of a relatively small subset of genes rather than large-scale genome restructuring. These results provide the first high-quality genomic resource for Bga and offer new insights into the evolutionary mechanisms underlying host specialization in powdery mildew fungi.

## Introduction

*Blumeria graminis* (formerly *Erysiphe graminis*) is an obligate biotrophic fungal pathogen responsible for powdery mildew diseases in cereals and grasses worldwide. This pathogen generally exhibits strict host specialization based on phenotypic criteria, with infection typically limited to a single host type, as first documented in several studies from the early-to-mid 20^th^ Century (Hardison 1945; Mains 1934; Marchal 1902; Oku et al. 1985). The economically important cereal-attacking forms of *B. graminis* are *B. graminis* f. sp. *tritici* (*Bgt*) infecting wheat, *B. graminis.* f. sp. *hordei* (*Bgh*) infecting barley, *B. graminis* f. sp. *secalis* (*Bgs*) infecting rye and *B. graminis* f. sp. *avenae* (*Bga*) infecting oat (Troch et al. 2012; Wyand and Brown 2003). Although several formae speciales can occasionally infect accessory hosts under experimental conditions, *Bga* is considered one of the most host-specialized members of the species complex, with infection largely restricted to species of *Avena* (Inuma et al. 2007; Wyand and Brown 2003).

Oat (*Avena sativa* L.) has attracted renewed interest in recent years owing to its nutritional value and increasing demand for healthy and plant-based food products. However, powdery mildew remains one of the most widespread diseases affecting oat cultivation. Severe infections reduce photosynthetic efficiency, accelerate leaf senescence and can lead to substantial losses in yield and grain quality (Jones 1977; Roderick et al. 2000; Różewicz et al. 2021). The disease is controlled primarily through the deployment of resistant cultivars, making knowledge of pathogen diversity and evolutionary potential crucial for effective breeding programs. Recent studies have demonstrated considerable diversity in virulence patterns among *Bga* populations and documented temporal changes in the effectiveness of resistance genes deployed in oat cultivars (Cieplak et al. 2022, 2021; Grzelak et al. 2025; Okoń et al. 2021). These observations suggest that *Bga* populations are continuously evolving in response to host selection pressure.

Understanding the molecular basis of virulence and avirulence in *Bga* is an essential requisite to elucidating mechanisms of resistance and adaptation in the *Avena*: *Bga* pathosystem. While several powdery mildew resistance genes have been identified in oat, little is known about the corresponding pathogen determinants recognized during host–pathogen interactions (Ociepa et al. 2020; Ociepa and Okoń 2022; Okoń 2015).

Phylogenetic studies based on individual loci, including ITS, 28S rDNA, β-tubulin and chitin synthase genes (Inuma et al. 2007; Wyand and Brown 2003) together with analyses of avirulence (AVR)-effector gene homologues (Sacristán et al. 2009) and cell wall penetration patterns (Tosa et al. 2011), have confirmed the distinct position of *Bga* among cereal powdery mildew pathogens and revealed genetic differentiation associated with geographic origin. However, analyses based on a limited number of molecular markers provide only a partial view of evolutionary relationships and cannot fully explain the mechanisms underlying host specialization and pathogenic adaptation. Recent genomic studies have demonstrated that adaptation in powdery mildew fungi is driven by variation in effector repertoires, transposable element (TE)-rich regions and structural rearrangements that cannot be captured using a small number of loci (Frantzeskakis et al. 2018; Müller et al. 2019), highlighting the need for whole-genome approaches to identify the genetic basis of divergence, host adaptation and virulence evolution (Badet and Croll 2020; Menardo et al. 2017).

Advances in next-generation sequencing have greatly improved our understanding of genome evolution in obligate biotrophic fungi. High-quality genome assemblies of *B. graminis* f. sp. *hordei* and *B. graminis* f. sp. *tritici* revealed large, repeat-rich genomes shaped by extensive TE proliferation and substantial structural plasticity (Frantzeskakis et al. 2018; Müller et al. 2019; Spanu et al. 2010). These features contribute to the rapid evolution of pathogenicity-related genes, particularly effectors, which are often located in repeat-rich regions and display elevated sequence diversification, copy number variation and presence–absence polymorphisms. Functional characterization of avirulence genes in wheat powdery mildew has demonstrated that variation in individual effectors can determine compatibility with host resistance genes and drive the emergence of new virulence phenotypes (Bourras et al. 2019; Praz et al. 2017).Consequently, characterization of the *Bga* effector repertoire represents an important step toward understanding the molecular mechanisms governing pathogenicity and host specificity in the *Avena*: *Bga* pathosystem.

Recent genomic studies have highlighted the importance of structural variation, including chromosomal rearrangements, gene duplications, deletions and TE insertions, as major sources of adaptive diversity in fungal pathogens (Badet et al. 2020; Frantzeskakis et al. 2018; Müller et al. 2019). However, the highly repetitive nature of powdery mildew genomes poses challenges for genome assembly and annotation. The integration of long-read sequencing, transcriptomics and Hi-C technologies has enabled chromosome-scale genome assemblies and improved gene annotation, facilitating increasingly detailed investigations of genome organization, effector evolution and host adaptation, as well as comparative and pangenomic analyses of fungal pathogens (Müller et al. 2018; Xia et al. 2022; Yildirir et al. 2022). These advances have enabled increasingly detailed investigations of genome organization, effector evolution and host adaptation, while also providing a foundation for comparative and pangenomic analyses of fungal pathogens (Badet and Croll 2020).

Despite the economic importance of oat powdery mildew, genomic resources for *Bga* remain scarce compared with those available for wheat and barley powdery mildew pathogens, for which chromosome-scale genome assemblies are already available. The absence of a high-quality reference genome has limited investigations of genome organization, repeat composition, effector evolution and the molecular basis of host specialization, while preventing comprehensive comparative analyses across cereal-infecting formae speciales. Closing this knowledge gap is essential for advancing our understanding of the evolutionary processes shaping pathogenicity in powdery mildew fungi.

Therefore, in this study, we undertook the first comprehensive genomic analysis of *Bga*. De novo assembly was generated using a combination of long-read PacBio (Sequel II) sequencing and paired-end Illumina HiSeq-PE150 sequence data, supplemented with transcriptomic analyses and Hi-C data to enhance assembly contiguity and gene annotation accuracy. We re-sequenced five *Bga* isolates with distinct virulence/avirulence profiles to capture the genetic diversity associated with host adaptation. This multi-platform genomic framework establishes the first high-quality reference genome for *Bga*, opening new research avenues for this pathogen similar to those enabled by the reference genomes of *Bgt* and *Bgh*. The availability of comprehensive genomic resources will facilitate advanced comparative analyses across *B. graminis* formae speciales, enable detailed studies of host specialization mechanisms, and provide a foundation for future investigations into pathogen evolution, effector biology, and host-pathogen interactions in the *Avena*: *Bga* pathosystem.

## Results

### Genome assembly

After filtering, PacBio long-read sequencing yielded 2,057,316 subreads with a total size of 32 Gb, average read length of 13,833 bp and NG_50_ of 15,843 bp. The estimated genome size of *Bga* from sub-sequences (k-mers = 21) of long-read data is approximately 139LMb, similar to that of other *B. graminis* species reported previously (Müller et al. 2019; Spanu et al. 2010). The total bases sequenced thus correspond to an average genome coverage of approximately 230x. The 21 k-mer spectrum of the long reads fits well with an expected haploid genome model (84.1%), where the heterozygosity level was very low with a mean rate of 0.15%.

The proportion of repetitive content, on the other hand, was substantial at ∼90 Mb. This level of repetitive content is comparable to *Bgt*, which comprises ∼90% of the genome (∼149LMb), and notably higher than *Bgh*, whose genome contains ∼64% repetitive content (∼80LMb). These findings highlight the extensive genome plasticity among *B. graminis* formae speciales, likely reflecting their adaptation to different hosts. The percentage of duplicated genes in our assembly was also lower than that of the previous draft genomes, with only 1.7% of duplicated BUSCO genes (Supplementary Table 1).

We generated separate assemblies from these reads using the PacBio assemblers FALCON and Canu, respectively. We selected the FALCON assembly for scaffolding and downstream analyses because it produced the most contiguous assembly for our *Bga* isolate, consisting of only 250 contigs and a smaller total genome size (136.6LMb) compared to the assembly generated using Canu (Supplementary Table 1).

With the assistance of high-throughput chromosome conformation capture (Hi-C) data for contig joining and Illumina short-read sequences for polishing, we generated a highly contiguous assembly of *Bga*, which consists of 11 pseudomolecules (i.e. chromosomes) capped by identified telomeric repeats (TTAGGG/CCCTAA) on both ends suggesting their near-complete status. Unplaced contigs were concatenated to a pseudo-scaffold with a ∼2.1Mb size of sequence. The sizes of the 11 chromosomes range from 6.6 to 20 Mb (Table 1, Supplementary Table 1) and contain only 10 sequence gaps.

**Table 1.**
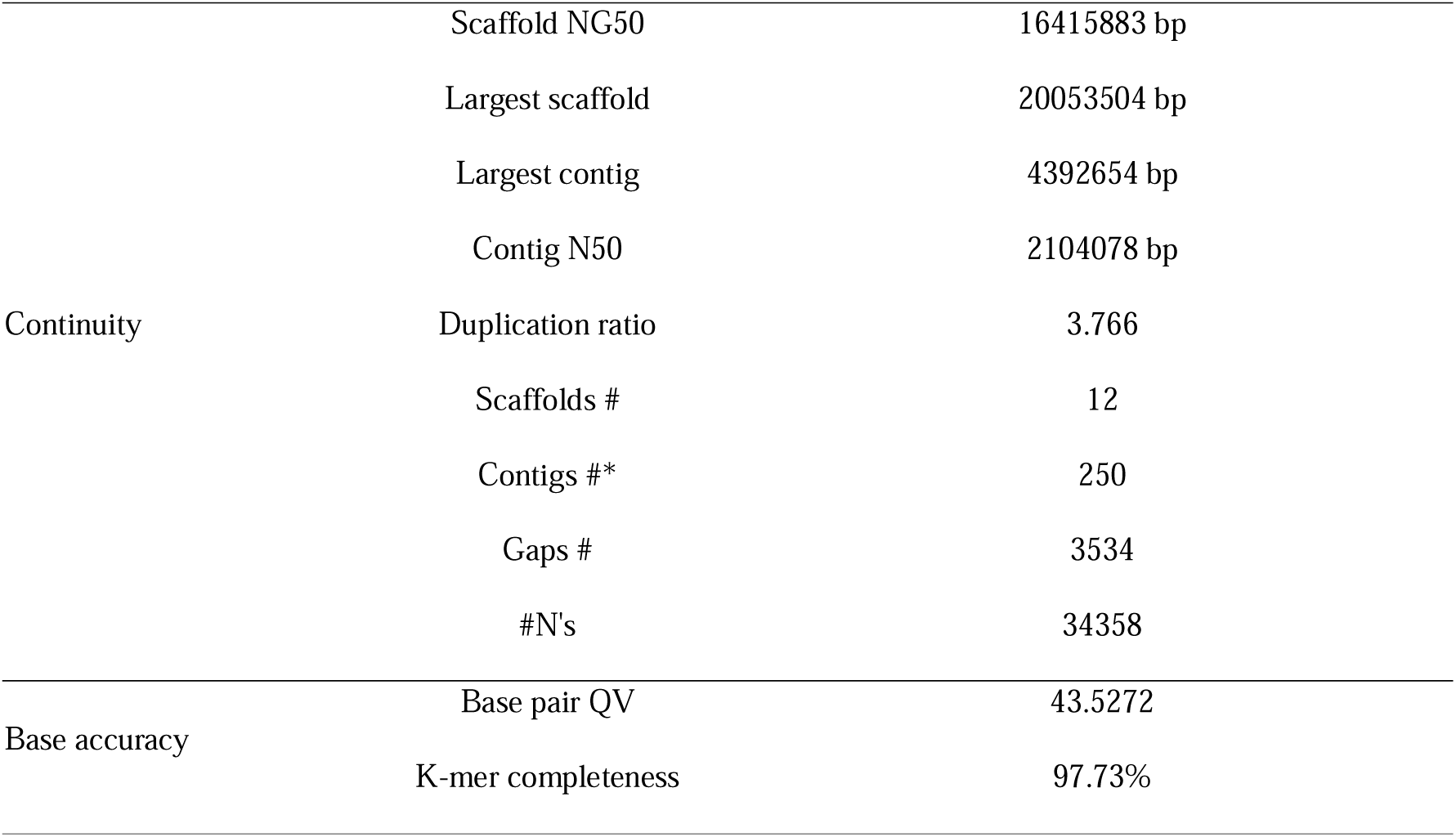
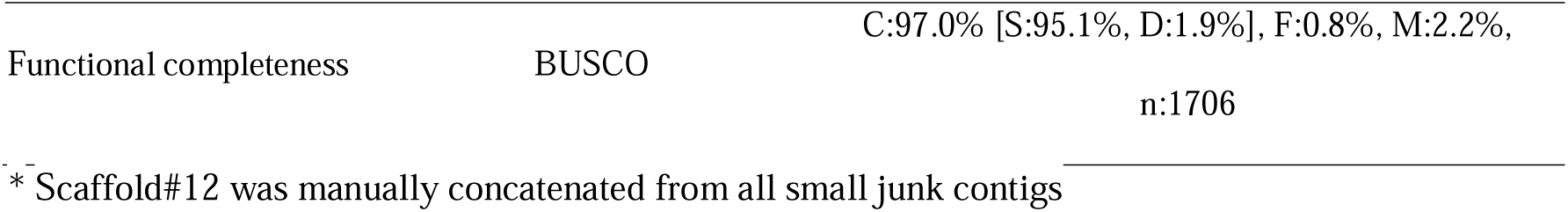
Assembly statistics for *Blumeria graminis* f. sp. *avenae* isolate 1D, pathotype TBPG.

By mapping genomic Illumina reads from five *Bga* isolates collected from diverse geographic locations (Table.L2) onto the reference assembly, we observed an overall even sequence coverage across all chromosomes, indicating a promising chromosome sequence continuity and consistency in these isolates. Chromosome size, GC content and repeat-rich regions also strongly correlated with gene content (Fig. 1). Different retrotransposon elements (RE) were evenly distributed along chromosome arms and interspersed with genes. A majority of highly condensed AT-rich/low GC content regions were mainly associated with long interspersed nuclear elements (LINEs) of the I/Jockey/L2 families, which were often associated with centromeres spanning a window size of ∼0.5-2 Mb (Fig. 1A, Fig. S1). These tandem repeat enriched regions aligned with higher-level symmetry and a layered organization of centromeric sequences (Fig. S1). The total interspersed repeats comprised a high percentage of the whole genome, occupying nearly 80%, underscoring the importance of repetitive DNA in shaping genomic architecture, with a portion of ∼22% still being unclassified (Supplementary Table 2).

**Figure 1.**
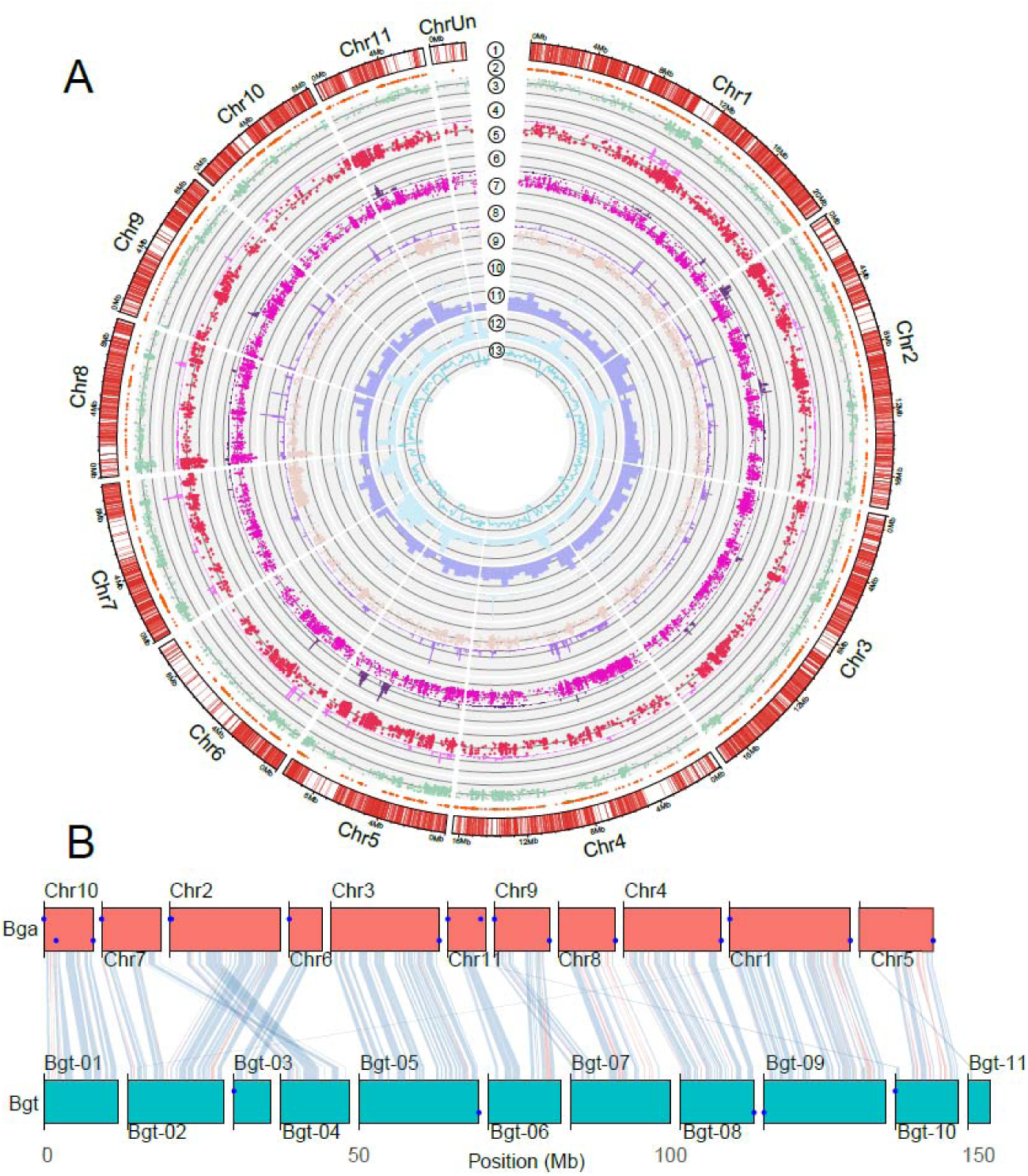
Genome organization and comparative synteny of *Blumeria graminis* f. sp. *avenae* (*Bga*). (A) Circular representation of genome-wide features across chromosomes of *Bga*. Tracks are shown from outer to inner as follows: (1) gene coordinates showing gene distribution along chromosomes; (2) conserved SNPs shared among four isolates; (3, 5, 7 and 9) SNP density for the four different *Bga* isolates with contrasting pathotypes, highlighting patterns of sequence variation; (4, 6, 8 and 10) isolate-specific homozygous SNP positions; (11) distribution of LINE-type transposable elements; (12) distribution of LTR-type transposable elements; and (13) GC content plotted as a continuous track. (B) Genome-wide macrosynteny between *Bga* (upper) and *B. graminis* f. sp. *tritici* (*Bgt*; lower). Chromosomes are represented as linear blocks, and collinear regions are connected by links, illustrating conserved genomic segments between species represented by BUSCO genes. The density and crossing of links reflect the extent of chromosomal rearrangements and conservation of gene order. Telomeric regions are indicated by dots at chromosome ends.

Macrosynteny analysis between *Bga* and *Bgt* revealed extensive conservation of chromosomal organization, with most chromosomes showing one-to-one correspondence and largely collinear gene order (Fig. 1B). Syntenic blocks spanned large chromosomal segments, indicating a high level of genome conservation between the two formae speciales. However, localized rearrangements, including inversions and translocations, were observed.

### The contribution of TEs to *Bga* genome structure based on long-term and rapid evolution

TEs can comprise large proportions of powdery mildew genomes, up to around 90% (Wicker et al. 2013). In *Bga*, repeat elements accounted for 80.48% of the whole genome, of which 51.86% were retrotransposons (24.86% and 27.01% were LINEs and long terminal repeat (LTR) elements, respectively), 5.69% were DNA transposons, and 22.44% were unclassified interspersed (Supplementary Table 2). TE evolution in powdery mildew fungi has been suggested to occur over both long-term and contemporary timescales, however, we found that TE sequences are divergent and that there were no obvious recent or ongoing TE bursts (Fig. 2A, 2B). We detected two weak peaks of contrasting TE composition, suggesting two TE burst events in the evolution of the *Bga* genome (Fig. 2A). To explore TE contributions, we examined where TE segments and exons overlapped in several genomic contents and regions. We found that the relative contribution of TEs to the different categories of genic features differed from a random model of overlap based on the frequency and coverage of TEs in the genome (Fig. 2C). TE contribution to functional genic features was greater for predicted small-secreted protein (SP) genes than for other gene types such as carbohydrate enzyme (dbCAN) genes but was still less than expected based on their sheer genomic abundance (Fig. 2C). A similar approach was used to evaluate TE contribution to other genomic elements, including AT-rich and topologically associating domain (TAD) boundary regions. TE contribution to the AT-rich regions was much stronger than to the TAD regions, with LTR being the most predominant (Fig. 2D).

**Figure 2.**
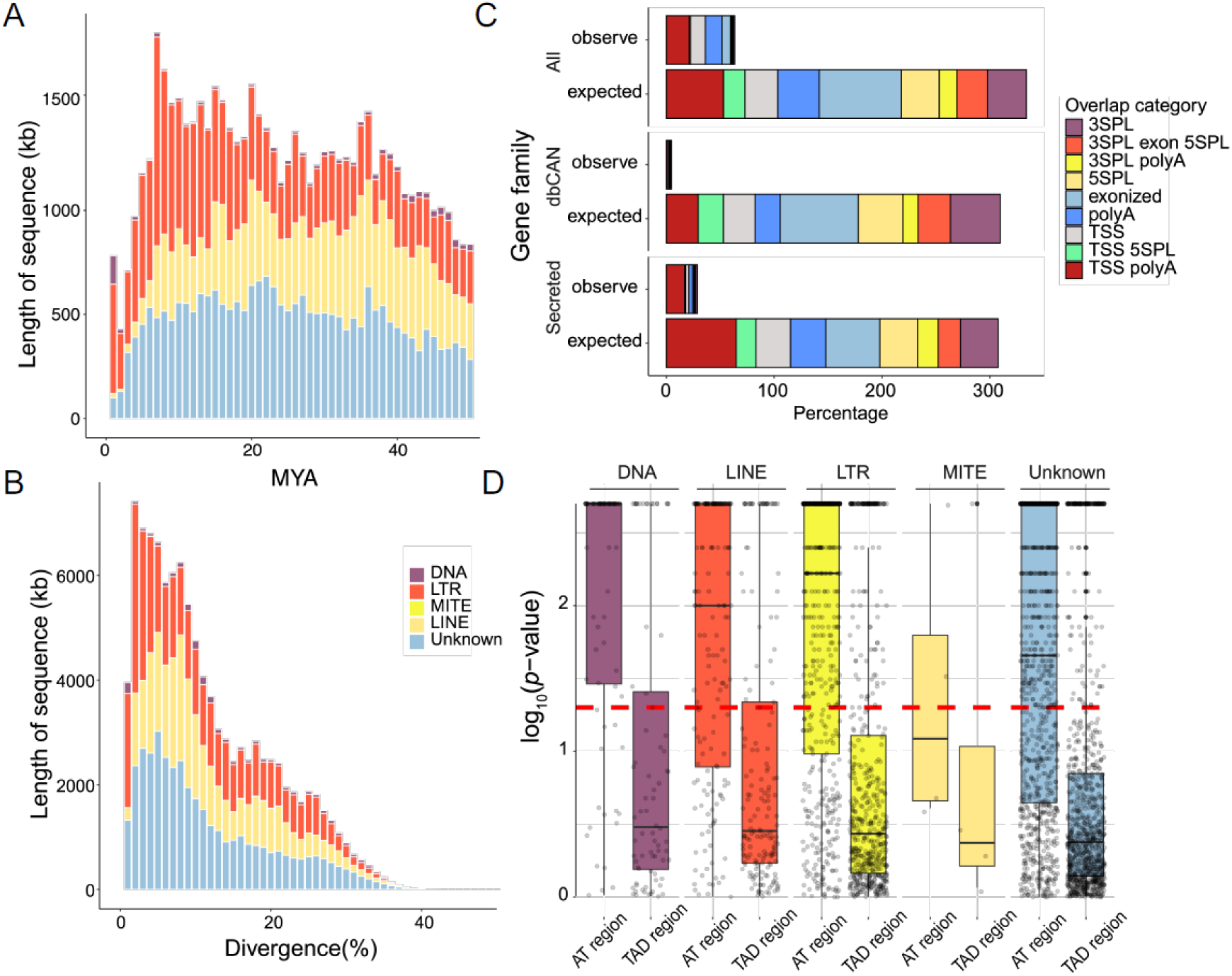
Contribution of transposable element (TE) to genomic features in *Bga*. (A) Temporal accumulation of TEs based on divergence estimates, showing the relative contribution of four TE classes (DNA, LTR, LINE, and unknown) across evolutionary time [million years ago (MYA)]. (B) Distribution of TE sequence divergence, indicating the abundance of TE classes across divergence bins. (C) Proportion of TE overlap across gene feature categories. Observed TE contributions are compared to expected values derived from randomized models accounting for genome-wide TE frequency and coverage. Overlap categories include transcription start site (TSS), splice site (SPL), exon-associated regions (TSS+SPL, both SPL), and polyadenylation-associated regions (polyA). Analyses are shown for all genes, carbohydrate enzyme (dbCAN) genes, and predicted secreted protein genes. (D) Enrichment of TE classes (DNA, LINE, LTR, MITE, and unknown) in different genomic regions. Distributions of log_10_(*p-value*) are shown for AT-rich regions and topologically associating domain (TAD) boundary regions, with significance thresholds indicated by dashed lines. Points represent individual genomic windows, and boxplots summarize enrichment distributions for each TE class and genomic category.

Many AT-rich sequences in fungal genomes are derived from the action of repeat-induced point-mutation (RIP) (John Clutterbuck 2011; Galagan and Selker 2004). The GC nucleotides in duplicated or repeated sequences were heavily mutated to contain numerous AT transition mutations to inactivate TEs and increase the local AT-content. Like all analysed powdery mildew fungi so far, RIP is also absent in the *Bga* genome. We found that AT-rich regions comprised ∼11% of the genome, and while they were interspersed in the *Bga* chromosomes they were located primarily within repetitive elements, especially LTR type repeat elements (Fig. 1A). These LTR elements contain long, repeated sequences at their ends and could be involved in the movement of genetic material within the genome. To further characterize the relationship between TEs, sequence composition, and gene organization, we examined dinucleotide bias, gene-TE distances, and intergenic structure across the genome (Fig. S2). Dinucleotide frequency analyses revealed clear differences among TE families, with ERVK elements exhibiting higher (CpA + TpG)/(ApC + GpT) ratios, indicative of stronger transition-associated mutation bias, whereas Gypsy elements showed elevated TpA/ApT ratios, consistent with their enrichment in AT-rich regions (Figs S2A-B). Analysis of gene-TE distances showed non-random spatial organization, with distinct upstream and downstream distance distributions and differences between gene categories, including secreted and non-secreted genes (Fig. S2C). Genome-wide intergenic distance analysis further revealed a bimodal distribution of gene spacing, reflecting the coexistence of gene-dense regions with short intergenic distances and gene-sparse regions with expanded intergenic space (Fig. S2D). Notably, putative secreted protein genes were preferentially located in regions with larger flanking intergenic distances, which are typically enriched in repetitive elements (Fig. 1A, Fig. S2D). Together, these patterns indicate that TE family composition, mutation signatures, and gene positioning are tightly associated, supporting a compartmentalized genome structure characterized by TE-rich, gene-sparse regions and gene-dense regions.

### Chromosomal structure analysis

Contacts between chromosomes were dominated by centromere-centromere contacts and contacts between subtelomeric regions (Figs 3A-C). This effect was the strongest for the AT-rich blocks, with contacts between these regions accounting for nearly half of all inter-chromosomal contact (Fig. 3D). These results suggest that the AT-rich blocks are key determinants of 3D genome structure in *Bga*, and that the genome is partitioned into AT-rich and gene-rich blocks in three dimensions as well as in one dimension. Like many other fungal species, we found minimal intrachromosomal compartmentations (Fig. 3B). Major compartmentalization was only found for heterochromatic centromeres and telomeres (Figs 3B, E). This phenomenon has been postulated as due to the integration of smaller heterochromatic regions into larger euchromatin domains (Xia et al. 2022), such as the structure of TAD-like regions. In filamentous fungi, TAD-like structures generally form through hierarchical clustering of local chromatin structures that are also known as regional globule clusters (Torres et al. 2023). TAD-like domains appear as triangular regions of increased contact frequency along the diagonal of Hi-C interaction matrices, which visualize contact frequencies between pairs of genomic loci. We found that predicted strong boundaries were predominantly located at the borders of AT-rich regions (Fig. S3), suggesting an association between sequence composition and local chromatin organization in Bga.

**Figure 3.**
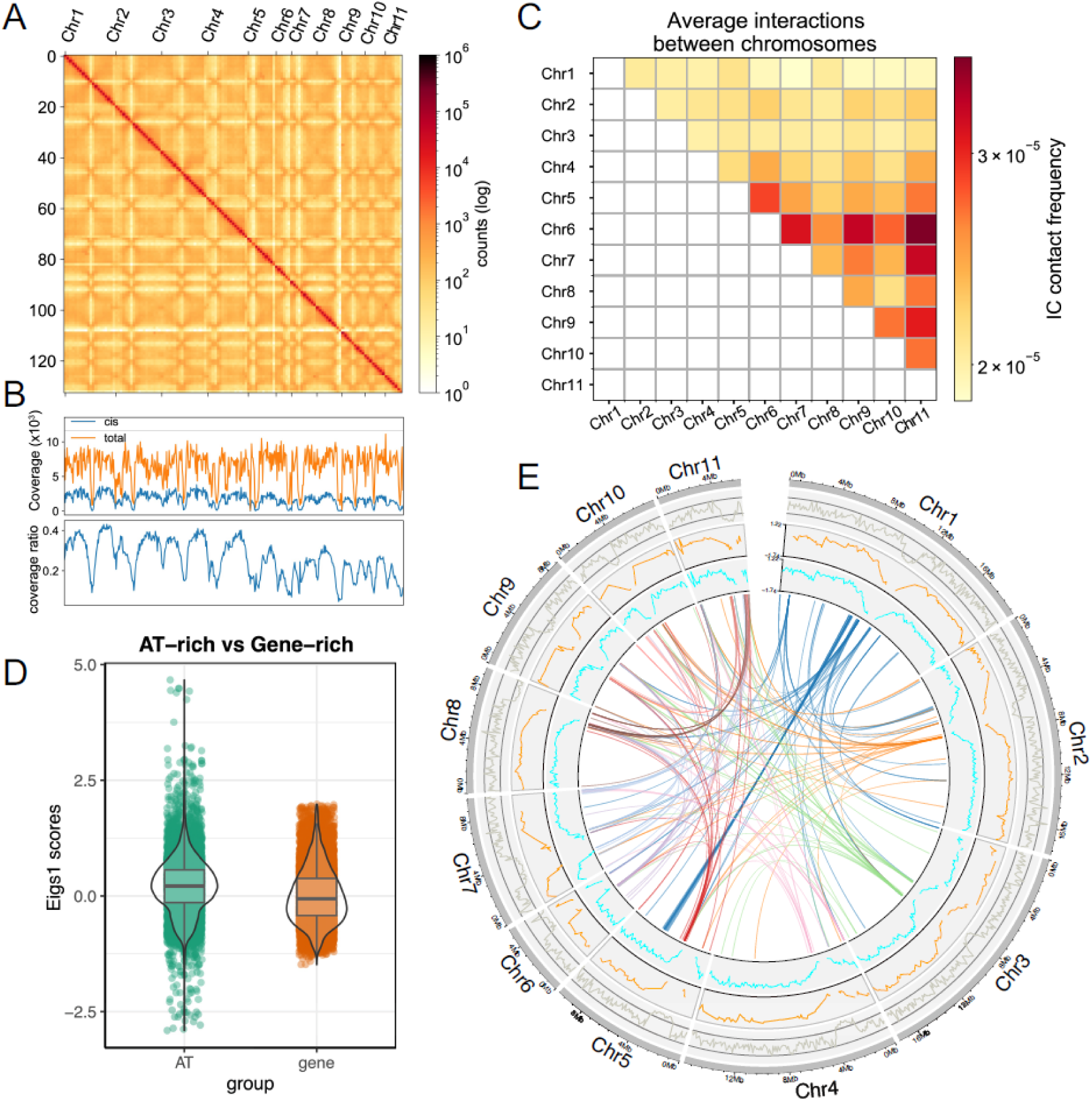
Chromosome conformation and compartmentalization of the *Bga* genome. (A) Genome-wide Hi-C contact matrix showing interaction frequencies between genomic regions across concatenated chromosomes. Both axes represent genomic coordinates (Mb), with chromosomes arranged sequentially. Interaction frequencies are displayed as log-transformed contact counts derived from the ICE-normalized matrix, with warmer colors indicating higher contact frequencies. (B) Summary of Hi-C interaction profiles across the genome. The upper panel shows interaction coverage (Hi-C contact counts per genomic bin) for total-and cis-interactions, and the lower panel shows the cis-to-total interaction ratio along genomic coordinates. (C) Average inter-chromosomal interaction frequencies between chromosomes. Heatmap values represent normalized contact frequencies from the balanced Hi-C matrix, with both axes indicating chromosome identities. (D) Distribution of first eigenvector (E1) scores derived from eigenvector decomposition of the normalized Hi-C contact matrix. E1 values are shown for genomic bins (100 kb) classified as AT-rich or gene-rich regions. Points represent individual bins, and violin plots summarize the distribution of E1 scores for each group. Differences between AT-rich and gene-rich regions were assessed using the Welch’s *t*-test, indicating a significant difference between groups (*t* = 25.09, *p* < 1 × 10^-130^). Bayesian analysis supported this result, with a large Bayes factor and a posterior difference estimate with narrow 95% credible intervals. (E) Circos representation of genome-wide chromatin interactions. Outer tracks show chromosomal coordinates and associated genomic features, including GC content and E1 profiles of gene-rich (orange) and AT-rich (cyan) regions. Links represent inter-chromosomal contacts between genomic regions.

Genome compartmentalization, referred as the two-speed genome, is often found in plant-pathogenic fungi and oomycetes (Dong et al. 2015). In two-speed genomes, compartments containing core-conserved genes (i.e. those involving housekeeping, metabolism, and development) are often gene-rich and repeat-sparse regions, while a substantial portion of the genome comprises enriched repeat elements likely driving virulence gene variation (Frantzeskakis et al. 2019; Raffaele and Kamoun 2012). On the contrary, powdery mildew fungi generally possess one-speed genomes in which drastic repeat-rich compartments are not found (Kusch et al. 2024). Like other characterised powdery mildew fungi, TEs in the *Bga* genome also contribute minimally to gene compartments (Fig. 2C).

### Gene content analysis

To investigate genomic contexts in terms of host and species-specificity, we compared global syntenic relationships for gene content among *Bga* and 16 powdery mildew fungi with either monocot or dicot hosts. Dynamics and properties of the entire synteny networks were implemented to identify patterns of genome evolution that could provide insights into how genome dynamics and gene content could potentially contribute to trait evolution (Zhao and Schranz 2019). The patterns of gene copy number across species of all clusters indicated that conserved single-copy syntenic clusters are predominant in the network (Fig. 4A). Overall, shared syntenic gene contents comprised 910 clusters with a total of 9,186 genes in the evaluated powdery mildew genomes. These genes were enriched for terms predominantly associated with macromolecule modifications (FDR 1.9e-06) such as the binding of nucleoside phosphate, carbohydrate derivatives, RNA and proteins, and with cellular component biogenesis (FDR 1.7e-08) such as chromatin organisation, histone modification and gene expression, which are all highly conserved pathways regardless of host range (Supplementary Table 3). Synteny of whole-genome protein sequences also revealed significantly more conserved gene clusters in the genomes of the monocot host clades, including *Bga*, *Bgt*, and *Bgh* (Fig. 4A). These monocot-specific gene contents fell into 904 clusters, in which grass-infection specificity of these pathogens was clearly reflected by the overrepresentation of metabolic processes of organonitrogen (FDR 2.4e-05), organophosphate (FDR 5.4e-05), dicarboxylic acid (FDR 1.7e-04), as well as stress responses as those responding to hydrogen peroxide (FDR 2.7e-05), inorganic substance (FDR2.8e-03), osmotic stress (FDR 2e-03) and to salt stress (FDR 3.70e-05) and temperature stimulus (FDR 0.011) (Supplementary Table 4). We did not observe obvious carbohydrate metabolic processes that are typically essential for host cell penetration, consistent with the contraction of carbohydrate metabolism gene families in powdery mildew fungi (Liang et al. 2018).

**Figure 4.**
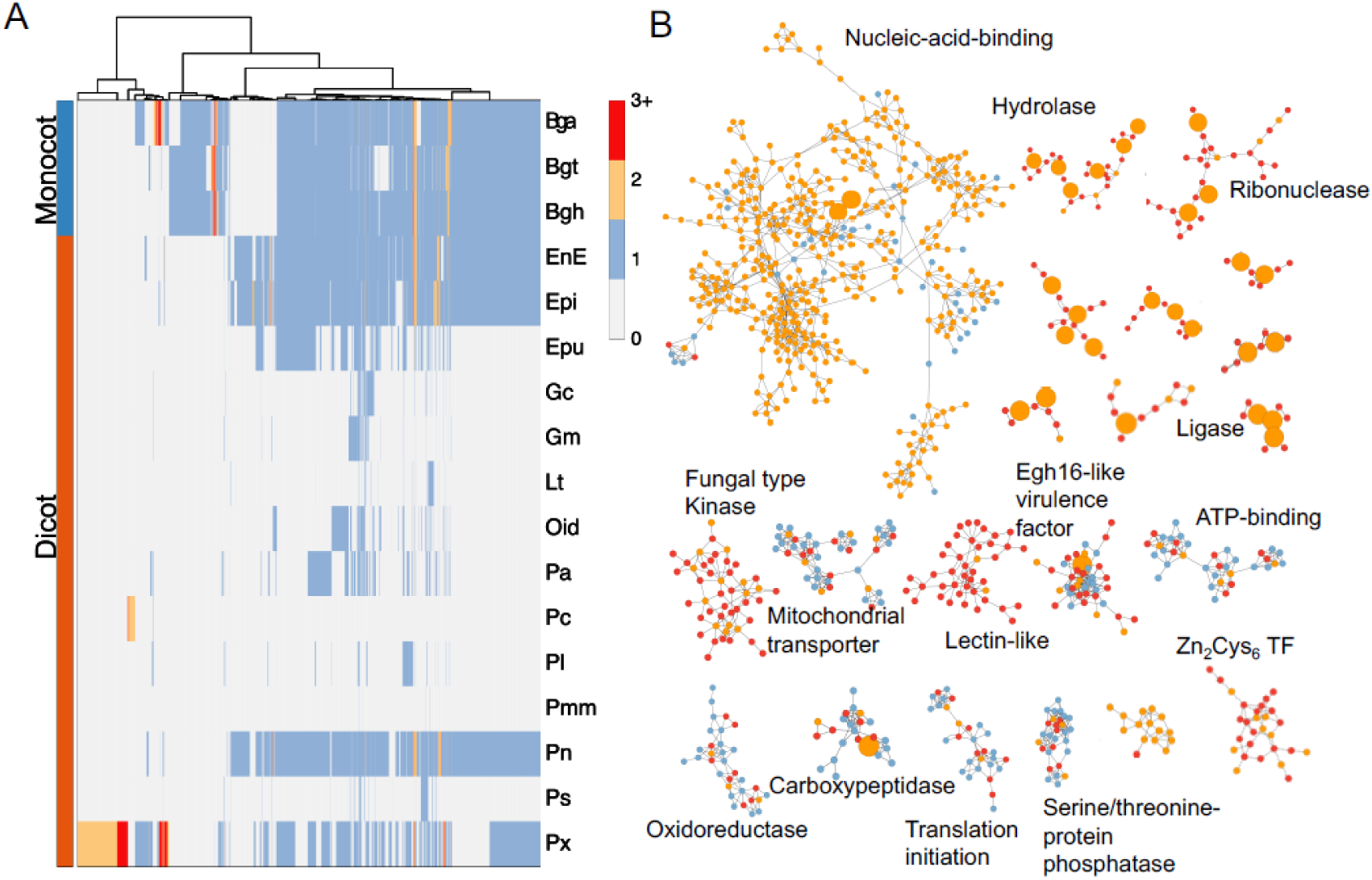
Global synteny relationships of gene content across *Bga* and 16 powdery mildew fungi. (A) Phylogenomic profile of synteny clusters showing patterns of gene copy number across *Bga* and 16 additional powdery mildew genomes. Columns represent synteny clusters and rows represent species. Values indicate the number of genes from each species within each cluster, with color scale reflecting copy number. Species are grouped according to host lineage (monocot and dicot powdery mildews), and hierarchical clustering is applied to both axes. (B) Representative synteny network clusters derived from the global synteny network. Nodes represent genes and edges represent syntenic relationships. Node color denotes copy number, and node size reflects the number of connections within the network. Selected clusters correspond to major functional categories, including nucleic acid-binding proteins, zinc finger transcription factors, kinases, mitochondrial transporters, lectin-like proteins, enzymatic proteins (e.g., ligases, hydrolases, ribonucleases), and other annotated functional groups. Networks illustrate both inter-and intra-species syntenic relationships across conserved gene families as well as clusters restricted to subsets of species. Bga, *Blumeria graminis* f. sp. *avenae*; Bgt, B. *graminis* f. sp. *tritici*; Bgh, *B. graminis* f. sp. *hordei*; EnE, *Erysiphe necator*; Epi, *E. pisi*; Epu, *E. pulchra*; Gc, *Golovinomyces cichoracearum*; Gm, *G. magnicellulatus*; Lt, *Leveillula taurica*; Oid, *Oidium heveae*; Pa, *Podosphaera aphanis*; Pc, *P. cerasi*; Pm, *Phyllactinia moricola*; Pmm, *Pseudoidium* sp.; Pl, *Podosphaera leucotricha*; Px, *P. xanthii*; Ps, *Pleochaeta shiraiana*.

Among the 2,594 genes of *Bga* that did not form any syntenic relationship, only about one third contain functional domains that are primarily associated with DNA/RNA binding. This group also included 164 predicted small secreted proteins, likely functionally specified as effectors for the oat host (Supplementary Table 5). We identified 322 *Bga*-unique syntenic genes in clusters of varying sizes, which are characterized predominantly by nucleic acid binding domains (Supplementary Table 6). These are likely due to homology protein domains or tandem repeat or protein fusions. We examined large-size clusters, which are likely maintained from several rounds of whole genome duplication events and/or tandem-duplicated arrays. These include DNA-binding domain containing gene group, zinc finger proteins, fungal-type kinases, enzymatic function proteins such as ligases, hydrolases, and ribonucleases, many of which are putatively secreted (Fig. 4B). In contrast, small clusters could represent lineage-specific transposition events, for which synteny is shared only across a few closely related species, such as transporter genes (Fig. 4B). Overall, the syntenic network reflects the differences and dynamics of conservation patterns among different genes and gene families in *Bga* along with other powdery mildew fungi.

### Transcriptome analysis during host infection

To investigate gene expression dynamics during host infection, RNA-seq analysis was performed on oat leaves inoculated with *Blumeria graminis* f. sp. *avenae* isolate 1D, representing pathotype TBPG. Samples were collected at 6, 24 and 48 hours post inoculation (hpi), and expression profiles were compared between the resistant genotype AV1860 (*Pm4*) and the susceptible cultivar Fuchs (Fig. 5). We examined three major functional gene categories: those encoding putative secreted proteins, transporter proteins, and carbohydrate-active enzymes. These gene groups did not show significant differences in overall expression between infections on resistant and susceptible host plants across timepoints (Figs 5A-C). Similar results were reported previously for the barley powdery mildew pathogen *Bgh* (Hacquard et al. 2013), suggesting that these core gene categories may be constitutively expressed during early infection stages. Despite a lack of clear global expression differences, DE analysis identified 58 and 345 genes that were significantly upregulated during infection of susceptible host at 24 and 48 hpi. At 24 hpi, corresponding to the early infection establishment stage, these DE genes are enriched GO terms that are mainly associated with cellular localization, particularly membranes and intracellular compartments, including the plasma membrane, organelle membranes, and organelles such as the nucleus, mitochondrion, vacuole, and Golgi apparatus (Fig. 5D).

**Figure 5.**
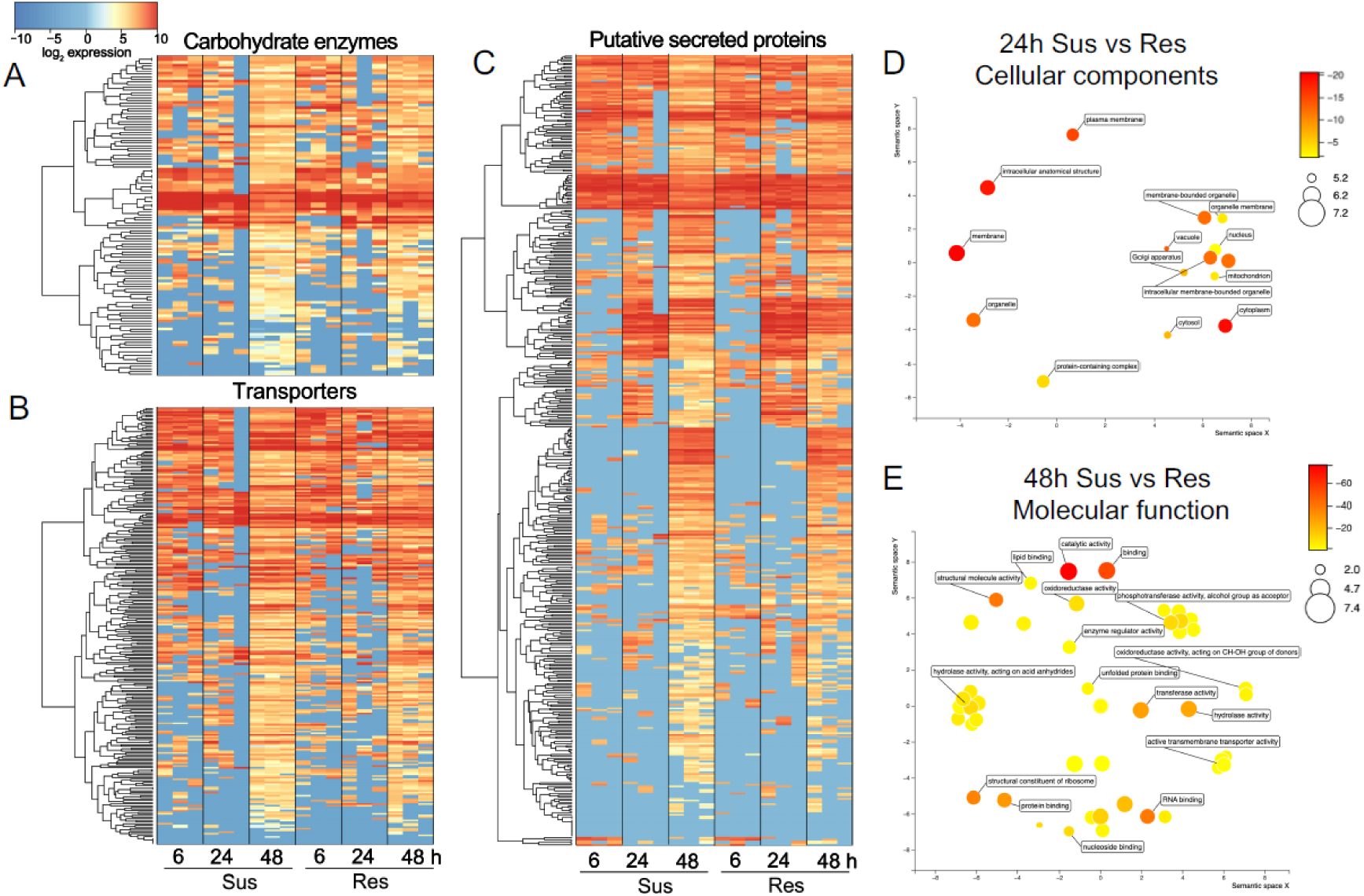
Gene expression dynamics in susceptible and resistant interactions. (A) Expression profiles of carbohydrate-active enzyme genes across susceptible and resistant host interactions at 6, 24, and 48 hours post inoculation (hpi). Heatmap showing expression values presented as loglJ-transformed FPKM values. Rows correspond to individual genes and columns to biological replicates for each condition and time point. Genes are ordered by hierarchical clustering based on expression similarity. (B) Expression profiles of transporter genes across susceptible and resistant conditions at 6, 24, and 48 hpi. (C) Expression profiles of putative secreted protein genes across susceptible and resistant conditions at 6, 24, and 48 hpi. (D) Gene Ontology enrichment of genes at 24 hpi for cellular component categories. Each node represents an enriched term, with node size proportional to the number of genes associated with the term and node color indicating enrichment significance (−log10 *p*-value). (E) Gene Ontology enrichment of genes at 48 hpi for molecular function categories.

By 48 hpi, gene enrichment had become more functionally diverse. GO terms related to catalytic and binding activities, including oxidoreductase, transferase, and hydrolase functions, as well as RNA/protein binding, ribosomal structural components, and active transmembrane transporter activity were all enriched, indicating broader metabolic and regulatory processes during later infection stages (Fig. 5E). Notably, many differentially expressed genes were located within genomic regions characterized by longer flanking intergenic regions and increased TE content (Fig. S4). This observation further supports the hypothesis that TE-rich genomic environments may facilitate the evolution of genes involved in host adaptation.

Because TADs represent structural domains of the genome where genes interact and can be co-regulated, examining the expression of genes located within these domains can reveal whether three-dimensional genome organization contributes to coordinated transcriptional responses during infection. Therefore, we investigated the expression patterns of genes located within TAD regions across infection stages and host interactions. Among the 1,145 genes located within TAD regions, only a small subset shows DE pattern during infection, but clear stage-and host-dependent patterns were documented. At 24 hpi, very few TAD-associated genes were differentially regulated, particularly in the susceptible interaction, where only six genes responded (Supplementary Table 7) indicating that transcriptional regulation within TADs is minimal during the early establishment phase. In contrast, the number of DE TAD genes had increased substantially by 48 h, especially in the susceptible host, where 86 genes were up-regulated compared with a more moderate increase in the resistant interaction (Supplementary Table 7). This suggests that transcriptional activation within TAD regions becomes more pronounced during later stages of infection and is associated with successful pathogen colonization. Functional annotations indicate that putative SPs are consistently represented among these genes, although they do not constitute the majority, while CANs are relatively rare but become more apparent at 48 h. Together, these results suggest that TAD-associated gene regulation intensifies as infection progresses and includes genes involved in host interaction, secretion, and metabolic processes during fungal growth in the susceptible host.

By intergenic distance analyses, we found that the *Bga* genome possesses gene contents displaying disparate flanking intergenic region (FIR) lengths (Fig. S4). The conserved syntenic genes across all powdery mildew species were significantly enriched in the short 3’ and 5’ FIR quadrant (Hg *P* = 1.32e-49, χ^2^ test *P* = 9.72e-55). On the contrary, genes unique to *Bga* were enriched in the long 3’ and 5’ FIR quadrant (Hg *P* = 7.51e-15, χ^2^ test *P* = 7.09e-15). Interestingly, differentially expressed genes tended to have longer 3’ and 5’ FIR. Among functional gene categories, only predicted effector genes showed significantly longer 3’ and 5’ FIR (Hg *P* = 6.16e-29, χ^2^ test *P* =5.16e-26). However, only 31 predicted effector genes were differentially regulated *in planta*, none of which showed synteny to genes in other powdery mildew fungi. *Bga* therefore tends to have a “one-speed” genome but with a pattern shifted towards faster evolution for host specific adaptation. Similar to the current finding is the gene-sparse pattern of candidate effectors displaying increased intergenic distances in the wheat powdery mildew fungus *Bgt*, indicating relatively “faster” evolution of host specific gene contents (Müller et al. 2019).

## Discussion

In this study, we present the first chromosome-scale reference genome of *Blumeria graminis* f. sp. *avenae* (*Bga*), providing a valuable resource for a *B. graminis* forma specialis that has remained largely unexplored despite its agricultural relevance. By integrating short-and long-read sequencing, Hi-C scaffolding and transcriptomics, we achieved a highly contiguous assembly that resolved key structural and functional features of the *Bga* genome. This resource substantially advances comparative genomic analyses within the *B. graminis* complex and provides new insight into the evolutionary mechanisms underlying host specialization.

The *Bga* genome exhibits hallmark features of powdery mildew fungi, including extensive repeat content and low gene density. The repeat fraction content (∼80%) places *Bga* closer to *Bgt* than *Bgh*, reinforcing the notion that genome expansion in this lineage is driven largely by transposable element proliferation rather than gene family expansion (Frantzeskakis et al. 2019; Müller et al. 2019; Spanu et al. 2010). TE-dominated genomes are increasingly recognised as important drivers of adaptive evolution in plant pathogens, where structural variation contributes to host adaptation and rapid evolutionary change (Badet and Croll 2020; Faino et al. 2016; Raffaele and Kamoun 2012). Unlike several filamentous fungal pathogens that exhibit clear signatures of recent TE bursts, our analyses suggest that TE activity in *Bga* is largely historical, with sequence divergence patterns consistent with ancient proliferation events. activity has been associated with increased genome plasticity and diversification of virulence-associated loci (Faino et al. 2016; Lorrain et al. 2021). The absence of strong recent TE expansion in *Bga* may indicate a relatively stable repeat landscape or, alternatively, suggest that adaptation has occurred through localized genomic restructuring. This observation suggests that TEs may have played a major role in the historical expansion of the *Bga* genome, whereas contemporary adaptation is more likely driven by local sequence diversification and regulatory changes affecting specific genomic regions. Such a scenario is consistent with the maintenance of a large, repeat-rich genome despite limited evidence for ongoing transpositional activity. We did not detect evidence for RIP, consistent with previous findings in powdery mildews (Frantzeskakis et al. 2019; Spanu et al. 2010), suggesting that long-term TE retention is a key factor shaping genome expansion in this lineage.

We found that *Bga* conforms to a “one-speed” genome model, as has been documented in other powdery mildew fungi (Kusch et al. 2024). Unlike the “two-speed” genomes observed in many filamentous pathogens, where gene-dense regions are separated from repeat-rich, rapidly evolving compartments (Dong et al. 2015), *Bga* displays a largely homogeneous genomic landscape with interspersed genes and repeats. Our data revealed that evolutionary heterogeneity is evident at a finer scale. Local genomic context, particularly FIR length, is strongly associated with gene function and evolutionary dynamics. Genes with long FIRs, including predicted effectors and differentially expressed genes, are enriched in TE-associated regions and exhibit reduced synteny conservation, whereas conserved genes are enriched in short FIR regions. This pattern indicates a functionally stratified “one-speed” genome, where evolutionary rates vary locally rather than across large compartments. Importantly, these findings indicate that adaptive evolution in *Bga* is concentrated within specific genomic neighbourhoods rather than within large accessory compartments. Thus, local genomic context appears to be a stronger determinant of evolutionary potential than chromosome-scale genome partitioning. Similar observations have been reported in other obligate biotrophs, where TE-rich environments contribute to regulatory plasticity and gene diversification without forming discrete genomic compartments (Duplessis et al. 2011; Frantzeskakis et al. 2019, 2018; Gupta et al. 2023).

Hi-C analysis provides further insight into genome organization, showing that AT-rich, repeat-dense regions play a central role in shaping three-dimensional chromatin architecture. Strong centromere-centromere and subtelomeric interactions are consistent with other plant associated fungi (Seidl et al. 2020; Winter et al. 2018). The identification of TAD-like structures, with boundaries enriched at transitions between AT-rich and gene-rich regions and depleted of TEs, suggests that genome folding is influenced more by sequence composition than by repeat density alone. Similar TAD-like domains have been described in filamentous fungi and are associated with chromatin organization and transcriptional regulation (Torres et al. 2023). In *Bga*, only a subset of TAD-associated genes is differentially expressed during infection, but the increase in transcriptional responsiveness at later stages particularly in susceptible interactions suggests that three-dimensional genome structure contributes to coordinated gene regulation during host colonization. Together, these observations indicate that chromatin architecture may represent an additional layer of regulatory control influencing infection-associated gene expression in *Bga*.

One of the central questions in *Blumeria* biology is how strict host specialization is maintained despite overall genomic similarity among formae speciales. Our results indicate that lineage-specific innovation in *Bga* is concentrated within a relatively small subset of genes. Predicted effectors are enriched among non-syntenic genes and TE-associated regions with extended FIRs, a pattern also reported in *Bgt* (Müller et al. 2019). These genomic contexts likely provide regulatory flexibility and promote sequence diversification. Collectively, these observations suggest that host specialization in *Bga* is driven by evolutionary changes affecting a relatively small subset of genes, particularly effectors and other infection-associated loci, rather than by extensive innovation across the entire genome. Recent structural studies of powdery mildew effectors have shown that many belong to conserved protein families with highly divergent sequences, enabling adaptation while maintaining functional constraints (Frantzeskakis et al. 2018; Müller et al. 2019; Praz et al. 2017). Notably, only a limited number of predicted effector genes are strongly differentially expressed during infection, suggesting that host specialization in *Bga* may depend on a combination of constitutive effector expression and fine-scale regulatory modulation rather than extensive effector expansion. The absence of strong CAN enzyme expansion further supports the notion that powdery mildew relies on intimate biotrophic interactions rather than on host tissue degradation (Hacquard et al. 2013; Liang et al. 2018). Instead, enrichment of genes involved in transport, signalling, and redox processes during infection indicates that metabolic adaptation and host manipulation are central to pathogenicity. Comparative synteny analyses highlight both conservation and divergence within the *B. graminis* complex. Conserved single-copy gene clusters dominate the genomic backbone and are enriched in essential cellular functions, including chromatin organization and gene expression, consistent with previous comparative studies (Frantzeskakis et al. 2019, 2018; Müller et al. 2019; Zaccaron et al. 2023). In contrast, *Bga*-specific and non-syntenic genes are enriched in predicted secreted proteins and nucleic acid-binding domains, suggesting roles in host adaptation. These findings support the hypothesis that host-driven selection acts primarily on a limited set of genes associated with host interaction, while the overall genomic framework remains highly conserved across cereal-infecting formae speciales. Specialization therefore appears to occur primarily through localized genetic and regulatory evolution rather than large-scale genome restructuring.

The availability of a high-quality *Bga* reference genome opens new opportunities for investigating the genetic basis of host specificity and virulence in the oat–powdery mildew pathosystem. In particular, population-scale resequencing of isolates with contrasting virulence profiles will be essential to identify candidate avirulence genes and to determine how genomic variation contributes to adaptation. Integration of such data with transcriptomic, epigenomic and functional analyses will help clarify how local genome organization, effector diversification and gene regulation interact during host colonization. These future approaches will be critical for linking genome architecture with pathogenicity and for translating genomic knowledge into more durable resistance strategies in oat breeding.

## Conclusions

This study provides the first chromosome-scale reference genome of *Blumeria graminis* f. sp. *avenae*, establishing an essential genomic resource for oat powdery mildew research. The *Bga* genome displays key genomic features of other powdery mildew fungi, including extensive repeat content, low gene density and the absence of clear evidence for recent transposable element expansion or RIP activity. These findings suggest that genome expansion in *Bga* has been largely shaped by historical TE proliferation and long-term repeat retention.

Our analyses indicate that *Bga* conforms to a functionally stratified “one-speed” genome model. Rather than being organized into large repeat-rich and gene-rich compartments, adaptive potential appears to be concentrated locally, particularly in TE-associated regions with extended flanking intergenic regions. These regions are enriched in predicted effectors, infection-responsive genes and non-syntenic loci, suggesting that local genomic context plays an important role in gene diversification, regulatory plasticity and pathogenic adaptation.

Comparative genomic and synteny analyses further show that host specialization in *Bga* is associated with localized genetic and regulatory changes affecting a limited subset of genes, while the overall genomic framework remains conserved across cereal-infecting formae speciales. Hi-C data additionally suggest that three-dimensional genome organization may contribute to the regulation of infection-associated genes during host colonization.

Together, these results provide new insights into the genome architecture and evolutionary mechanisms underlying host specialization in *Bga*. The reference genome presented here will support future population genomic analyses, identification of avirulence determinants, functional validation of candidate effectors and the development of more durable powdery mildew resistance strategies in oat breeding.

## Materials and methods

### *Bga* isolates

Five isolates of *Bga* were selected for this genomic investigation. The isolates were chosen to represent different levels of virulence, as determined by their differential responses to a panel of control genotypes harbouring known powdery mildew (*Pm*) resistance genes. Furthermore, these isolates were collected from distinct geographic regions to capture potential population-level genetic diversity. Detailed characterization of each isolate, including their geographic origin and virulence profiles against the differential host genotypes, is summarized in Table 2. The characterization is based on many years of observations conducted at the Institute of Genetics, Breeding and Biotechnology of Plants at the University of Life Sciences in Lublin (Cieplak et al. 2022; Okoń et al. 2021).

**Table 2.** Characteristics of *Bga* isolates used for PacBio and Illumina sequencing.

| Sample | Geographical origin | Isolate code | Reference genotype with different Pm genes |  |  |  |  |  |  |  |  |  |  |  |  |  |  |
| --- | --- | --- | --- | --- | --- | --- | --- | --- | --- | --- | --- | --- | --- | --- | --- | --- | --- |
|  |  |  | Pm1 | Pm2 | Pm3 | Pm4 | Pm5 | Pm6 | Pm7 (APR 122) | Pm7 (Canyon) | Pm3+8 | Pm9 | Pm10 | Pm11 | Pm12 | U | Fuchs (Susceptible) |
| 1D | Poland | TBPG | 4 | 0 | 4 | 0 | 0 | 4 | 1 | 3 | 4 | 1 | 4 | 4 | 0 | 4 | 4 |
| 2D | Poland | JBPB | 0 | 0 | 4 | 0 | 0 | 4 | 1 | 4 | 2 | 2 | 4 | 3 | 0 | 0 | 4 |
| 3D | Germany | TBDB | 4 | 0 | 4 | 0 | 0 | 4 | 0 | 2 | 4 | 2 | 3 | 2 | 1 | 0 | 4 |
| 4D | Ireland | TBRB | 4 | 0 | 4 | 0 | 0 | 4 | 1 | 3 | 3 | 3 | 2 | 3 | 0 | 1 | 4 |
| 5D | Poland | NBCB | 4 | 0 | 0 | 0 | 0 | 4 | 0 | 2 | 0 | 1 | 2 | 3 | 0 | 1 | 4 |

Isolate 1D with the highest virulence level was selected as the reference isolate for de novo genome assembly. The remaining diverse isolates were utilized for Illumina short-read sequencing. All isolates were propagated on leaves of the susceptible oat cultivar Fuchs. Spores were collected using a Mini Cyclone Spore Collector (Tallgrass Solutions, Inc.). DNA isolation was performed immediately after spore collection.

## DNA and RNA preparation for sequencing

### DNA extraction and quality control for PacBio long-read sequencing

Approximately 400 mg of spores were homogenized in liquid nitrogen using a pre-chilled mortar and pestle. DNA isolation from isolate 1D was performed according to the methodology described by Feechan et al. (Feehan et al. 2017), specifically optimized for long-read sequencing applications. This protocol ensures the extraction of high molecular weight (HMW) DNA crucial for generating long-read sequences and maintaining genome integrity during assembly. The purified genomic DNA was resuspended in TE buffer (10 mM Tris-HCl, 1 mM EDTA, pH 8.0) and stored at-20°C prior to shipment. DNA concentration and purity were determined using both spectrophotometric analysis with NanoDrop 2000 (Thermo Fisher Scientific) and fluorometric quantification using Qubit 4 (Thermo Fisher Scientific). DNA integrity was assessed by agarose gel electrophoresis, confirming that the majority of DNA fragments exceeded 20 kb in size.

### DNA extraction for Illumina short-read sequencing

Genomic DNA from isolates 2D-5D was extracted using a modified protocol based on Feechan et al. (Feehan et al. 2017), adapted for short-read sequencing requirements. For each isolate, approximately 150 mg of freshly harvested spores were ground to a fine powder in liquid nitrogen using a mortar and pestle. Key modifications to the original DNA isolation methodology included the incorporation of vortex mixing steps to enhance cell lysis and the termination of the extraction procedure at the first precipitation stage to obtain DNA suitable for Illumina sequencing. Following initial purification, samples were treated with RNase A to remove residual RNA contamination and further purified using AMPure XP magnetic beads (Beckman Coulter) according to the manufacturer’s protocol to ensure high-quality DNA recovery. The purified genomic DNA was finally resuspended in 22.5 μl of nuclease-free water. DNA concentration and purity were determined using NanoDrop 2000 and Qubit 4 (Thermo Fisher Scientific).

### RNA extraction for RNA-seq

Ten-week-old seedlings of the resistant oat genotype AV1860, carrying the *Pm4* powdery mildew resistance gene, and the susceptible cultivar Fuchs were inoculated with *Blumeria graminis* f. sp. *avenae* isolate 1D. Infected leaves were collected at 6-, 24-, and 48-hours post inoculation (hpi) in three replications. RNA was isolated using TRIzol reagent (Invitrogen) according to the manufacturer’s instructions.

### Hi-C library preparation

For chromosome-level scaffolding, Hi-C libraries were prepared from fresh conidia (approximately 1 g). Spores were resuspended in 1% formaldehyde solution (prepared from 37% stock in phosphate-buffered saline) at a 10:1 volume ratio and vacuum infiltrated for 30 min at room temperature with periodic vortexing. The crosslinking reaction was quenched by adding glycine to a final concentration of 125 mM, followed by an additional 15-minute incubation at room temperature with vortexing. The crosslinked material was pelleted by centrifugation (1000 × g, 1 min), washed with water, and re-pelleted. The crosslinked spores were then cryogenically ground to a fine powder in liquid nitrogen and transferred to clean tubes. The pelleted sample was shipped to Phase Genomics (Seattle, WA) for chromosome conformation capture sequencing.

### Whole-genome sequencing and reads processing

Prior to sequencing, all genomic DNA and RNA samples underwent quality control assessment using a Bioanalyzer (Agilent Technologies), with a minimum quality parameter threshold of 8 required for processing. Genomic DNA and RNA samples were sent to Novogene (Hong Kong, China) for Illumina short-read sequencing. TruSeq fungal libraries with 350bp insert were constructed, and 150bp paired-end (PE150) sequencing was performed on an Illumina NovaSeq 6000 platform, generating approximately 10Gb and 15Gb raw data for DNA-seq and RNA-seq, respectively.

PacBio long-read sequencing was performed on a single Sequel II cell using a circular consensus sequencing (CCS) library at Novogene, yielding approximately 80Gb raw sequence data. Hi-C sequencing (PE150) was performed at Phase Genomics (Seattle, WA). Genome assembly was performed following the PacBio Falcon (v1.8.1) (overlap_filtering_setting: --max-diff 200 --max-cov 600 --min-cov 3 --bestn 10; pa_DBsplit_option:-a-x500-s400; pa_HPCdaligner_option:-v-B128 - M32; pa_daligner_option:-k18-e0.75-l2500-s100-h1024-w8) and Canu (v1.9) (corMhapOptions=--threshold 0.80 --num-hashes 512 --num-min-matches 3 --ordered-sketch-size 1000 --ordered-kmer-size 14 --min-olap-length 1000 --repeat-idf-scale 50). For polishing of the assembly, three rounds of GCpp/Arrow (v2.0.1) and two rounds of pilon (v1.2) were used following Falcon.

### Annotation

Gene annotation was performed using an evidence-based pipeline integrating RNA-seq data, transcript assemblies, and protein homology. RNA-seq reads were aligned to the reference genome using HISAT2 (v2.2.1). Genome-guided transcript assemblies were generated using StringTie (v2.1.7), and *de novo* transcript assembly was performed using Trinity (v2.13.2). Assembled transcripts were processed with TransDecoder (v5.5.0) to predict coding sequences and generate protein-coding gene models. Transcript evidence was refined using the PASA pipeline (v2.5.2), which aligned transcripts to the genome using BLAT, GMAP, and minimap2 to produce high-confidence transcript structures and update gene models. Protein homology evidence was incorporated using ProtHint (v2.6.0) to generate extrinsic hints for gene prediction. *Ab initio* gene prediction was performed using BRAKER2 (v2.1.5), integrating RNA-seq alignments and protein-derived hints to train GeneMark-EX and AUGUSTUS models. Separate BRAKER2 runs were conducted using RNA-seq and protein evidence, and resulting predictions were combined using TSEBRA (v1.0.3) to generate a consensus gene set.

### Hi-C and chromatin structure analysis

Hi-C-based genome scaffolding was performed using the 3D-DNA pipeline in conjunction with the Juicer workflow (Durand et al. 2016). Raw Hi-C reads were processed, aligned, and filtered using Juicer, and scaffolding was conducted with 3D-DNA using default parameters. Processed Hi-C data were further handled using runHiC to generate remapped contact files and to convert outputs to multi-resolution Cooler (mcool) format for downstream analysis. Hi-C contact matrices were analyzed using the cooltools suite (Open2C et al. 2024). Contact matrices were generated at 100 kb resolution and normalized using iterative correction (ICE) to account for coverage and technical biases. Balanced contact maps were visualized as heatmaps to assess chromosomal interaction structure. Chromatin compartments were inferred by eigenvector decomposition of normalized contact matrices. The first eigenvector (E1) was computed for each chromosome, oriented based on gene density, and assigned to non-overlapping 100 kb genomic bins, where positive and negative values correspond to A and B compartments, respectively. To relate chromatin organization to genome features, GC content and gene density were calculated for each 100 kb bin from the reference genome sequence and gene annotation. E1 values were aligned with these features to assess associations between compartment identity and sequence composition. Genome-wide interaction patterns were visualized using heatmaps, genomic tracks, and circos-style plot with shinnyCircos (Wang et al. 2023).

### Gene synteny and synteny network analysis

Chromosomal syntenic analysis was performed using chromsyn (v1.2) (https://github.com/slimsuite/chromsyn). Comparative synteny analyses were performed using the syntenet R package, following the previously described syntenic network analysis pipeline (Zhao et al. 2021). Synteny inference in syntenet requires, for each genome, a set of translated protein sequences representing primary transcripts and a corresponding gene annotation file containing genomic coordinates. Accordingly, protein and gene annotation files were imported as species-wise lists of amino acid sequences and genomic ranges, respectively. Sequence identifiers were standardized to match gene identifiers in the annotation prior to downstream analysis. Pairwise sequence similarity searches were performed in an all-versus-all manner using DIAMOND BLASTp, and the resulting tabular similarity outputs were used as input for synteny detection. Syntenic blocks and anchor relationships were then inferred in syntenet using its native implementation of the MCScanX algorithm. The resulting synteny relationships were represented as an edge list, in which genes correspond to nodes and syntenic connections correspond to edges.

The inferred synteny network was clustered where each gene was assigned to a single synteny cluster, and cluster composition across species was summarized by phylogenomic profiling, in which rows represent synteny clusters and columns represent species. The resulting profile matrix records the copy number of genes from each species in each cluster and was used to identify deeply conserved clusters as well as lineage-restricted or genome-specific clusters. For downstream comparative analyses, genes were classified according to their participation in synteny clusters across 17 mildew genomes. Conserved genes were defined as genes belonging to clusters detected across a broader set of powdery mildew fungi, whereas non-syntenic genes were defined as genes lacking placement in conserved syntenic clusters. Genes without detectable homologs in *Bga* relative to the comparative mildew dataset were classified separately as genes without homologs. Cluster-based presence/absence patterns were further used to distinguish conserved versus lineage-restricted microsynteny relationships.

Synteny network visualization was based on the cluster assignments and corresponding phylogenomic profiles. Cluster-level heatmaps were generated from the phylogenomic profile matrix to display the distribution and abundance of syntenic genes across species. Network representations of selected clusters were constructed from the synteny edge list, with each node representing a gene and edges representing inferred syntenic links. Functional annotation classes associated with selected clusters were based on Phyre2 (https://www.sbg.bio.ic.ac.uk/phyre2/html/page.cgi?id=index) annotation and summarized for interpretation of cluster content. Molecular network was visualized in Cytoscape (v3.10.1).

### TE composition, enrichment, and genome-wide distribution

TEs were annotated using RepeatMasker (v4.1.5). The final genome assembly was screened against curated repeat libraries, and RepeatMasker was run with default sensitivity parameters to identify and classify interspersed repeats and low-complexity regions. RepeatMasker output files were retained for downstream analyses. TE annotations were extracted and genomic coordinates were parsed and converted into BED format (https://github.com/4ureliek/Parsing-RepeatMasker-Outputs). TE entries were filtered and reformatted to generate standardized annotation files, including TE coordinate tracks, TE class and family assignments, and genome-wide repeat coverage tables. Overlapping TE annotations were resolved by retaining primary annotations based on RepeatMasker scoring, and fragmented annotations belonging to the same TE instance were merged where applicable. Genome-wide TE coverage was calculated by summing the total length of annotated TE regions and normalizing by genome size. TE annotations were further grouped by class and superfamily to generate summary statistics for downstream analyses.

### TE divergence, enrichment and feature overlap analysis

TE composition and enrichment analyses were performed using the TE analysis pipeline (https://github.com/4ureliek/TEanalysis) (Kapusta et al. 2013). RepeatMasker-derived TE annotations were used as input, and genomic feature annotations were provided as comparison datasets. TE-feature overlaps were computed using BEDTools intersect, with parameters configured to report both partial and complete overlaps between TE regions and genomic features. TE content was quantified for each feature set, including total TE coverage, proportion of bases overlapping TEs, and distribution of TE classes and families.

TE landscape analyses were generated from RepeatMasker divergence estimates and summarized by TE class. In brief, TE coverage was computed as the proportion of bases annotated as TEs within defined genomic windows or feature sets. TE landscape analyses were generated using divergence estimates reported in RepeatMasker outputs, based on Kimura distance from consensus sequences. Divergence values were used to summarize TE age distributions across classes and families. TE landscape profiles were constructed by binning TE copies according to divergence values and calculating the relative contribution of each TE class across divergence bins.

TE insertions were assigned to functional overlap categories based on their positions relative to gene architecture. Categories were defined as follows: transcription start site (TSS), first exon including transcription start site and splice donor (TSS+SPL), splice site (SPL), internal exon including both splice sites (both SPL), polyadenylation site (polyA), and last exon including splice site and polyadenylation signal (polyA+SPL). TE insertions overlapping multiple features were assigned according to hierarchical classification rules to avoid multiple counting. To avoid redundancy from alternative splicing, a single representative transcript isoform per gene was retained. Only gene models containing complete structural annotations were included in the analysis. Observed counts of TE insertions within each overlap category were compared against a null distribution generated by randomizing TE coordinates across the genome. TE positions were shuffled within chromosomes while preserving chromosome lengths and excluding assembly gaps. Randomized datasets were generated across multiple iterations (n = 5,000), and overlap counts were recalculated for each iteration. Empirical distributions from randomized datasets were used to estimate expected values and to calculate enrichment statistics for each overlap category. TE enrichment analyses were additionally performed using permutation-based procedures implemented in the TE analysis workflow. Genomic feature coordinates were randomized while preserving chromosome identity, and observed TE overlaps were compared with randomized expectations across repeated iterations. Resulting overlap and enrichment statistics were compiled into summary tables for downstream visualization and statistical testing.

### Expression analysis

RNA-seq reads were aligned to the reference genome using HISAT2 (v2.2.1) with default parameters. Alignment files were sorted and indexed using SAMtools, and only uniquely mapped reads were retained for downstream analyses. Transcript assembly and quantification were performed using StringTie (v2.1.7) in a genome-guided manner. Transcripts were assembled independently for each sample, and a unified transcript annotation was generated by merging assemblies across all samples.

Expression levels were estimated as FPKM (Fragments Per Kilobase of transcript per Million mapped reads), and gene-level expression matrices were generated from StringTie outputs. Expression values were normalized across samples using the merged annotation to ensure consistency across conditions. Differential expression analysis was conducted using the Ballgown R package. Gene expression levels were compared between susceptible and resistant conditions at multiple time points (0, 24 and 48 hpi). Statistical significance of differential expression was assessed using Ballgown and genes were classified as differentially expressed (DE) based on model-derived *p*-values and log2 fold-change estimates. Resulting DE gene sets were used for downstream analyses, including integration with TE proximity, chromatin organization, and synteny-based gene classification.

Normalized expression values were used for downstream visualization, including heatmap representation of gene expression patterns across conditions and time points. Functional enrichment analyses of differentially expressed genes were performed based on gene annotations, and enriched GO functional categories were visualized using reviGO, in which node size reflects gene counts and node color represents enrichment significance.

### Gene-TE distance and intergenic region analysis

Spatial relationships between genes and TEs were quantified using the 2-speed genomes pipeline (https://github.com/rhysf/2speed_genomes) (Wacker et al. 2023). In brief, gene annotations and TE coordinates were used as input. Distances between genes and the nearest TE insertion were calculated using BEDTools-based nearest-neighbor approaches implemented within the pipeline. Distances were computed separately for upstream (5′) and downstream (3′) regions relative to gene boundaries, taking gene strand orientation into account. Overlapping features were retained as zero or negative distances according to pipeline defaults. Intergenic regions were defined based on adjacent gene coordinates, and intergenic distances were calculated as the genomic distance between neighboring gene boundaries.

Genes were classified into multiple categories for comparative analyses. DE genes were identified based on expression differences between susceptible and resistant host conditions. Conserved genes were defined as genes with orthologs present across 18 powdery mildew fungal genomes. Non-syntenic genes were defined as genes lacking conserved genomic position relative to orthologous loci across these genomes. Genes without homologs were defined as lineage-specific genes without detectable sequence similarity to genes in other powdery mildew fungi. For each gene category, distributions of gene-TE distances and intergenic distances were compared to genome-wide background distributions. Four groups portioned by 5′ and 3′ median intergenic distances were used to assess relationships between flanking intergenic region (FIR) lengths and gene functional and expressional categories. Gene numbers in each partitioned quadrants were tested for enrichment using hypergeometric (Hg) and chi-squared (χ^2^) tests.

## Declarations

Ethics approval and consent to participate: Not applicable

Consent for publication: Not applicable

## Availability of data and materials

The datasets generated and analysed during the current study are available in publicly accessible repositories. The reference genome assembly, genome annotation files, analysis scripts and additional supporting files are available in the GitHub repository: https://github.com/yiding1121/Pma-data/tree/main.

Raw sequencing data generated in this study, including PacBio long-read sequencing, Illumina short-read sequencing, Hi-C sequencing and RNA-seq data, have been deposited in the NCBI Sequence Read Archive under BioProject accession number PRJNA1492323.

Raw data underlying selected main figures are provided in the updated Supplementary Data file. Additional data generated or analysed during this study are included in this article and its supplementary information files.

## Acknowledgements

The authors gratefully acknowledge the generosity of Judith and Davind Coffey and family for financially supporting this research.

## Authors’ contributions

YD participated in project conception, data analyses and interpretation and manuscript drafting. PZ participated in project conception, sequencing facilitating and manuscript preparation.

TO participated in spore collection and genetic material isolation, and prepared the initial version of the manuscript.

AN participated in sample preparation, spore collection, and genetic material isolation. HG conducted initial transcriptome analyses.

KK participated in developing the research concept and experimental plan.

RP Conceptualized and initiated the project, secured funding, provided overall supervision and contributed to preparing the manuscript.

SO participated in developing the research concept and experimental plan. She was the main investigator during the spore collection and genetic material isolation stages and contributed to writing the manuscript.

